# Deep-Learning-Enabled Differentiation between Intraprostatic Gold Fiducial Markers and Calcification in Quantitative Susceptibility Mapping

**DOI:** 10.1101/2023.10.26.564293

**Authors:** Ashley Wilton Stewart, Jonathan Goodwin, Matthew Richardson, Simon Daniel Robinson, Kieran O’Brien, Jin Jin, Markus Barth, Steffen Bollmann

## Abstract

**Purpose:** Interest is growing in MR-only radiotherapy (RT) planning for prostate cancer (PCa) due to the potential reductions in cost and patient exposure to radiation, and a more streamlined workflow and patient imaging pathway. However, in MRI, the gold fiducial markers (FMs) used for target localization appear as signal voids, complicating differentiation from other void sources such as calcifications and bleeds. This work investigates using Quantitative Susceptibility Mapping (QSM), an MRI phase post-processing technique, to aid in the differentiation task. It also presents deep learning models that capture nuanced information and automate the segmentation task, facilitating a streamlined approach to MR-only RT.

**Methods:** CT and MRI, including GRE and T1-weighted imaging, were acquired from 26 PCa patients, each with three implanted gold FMs. GRE data were post-processed into QSM, *T* 2^*^, and *R*2^*^ maps using QSMxT’s body imaging pipeline. Statistical analyses were conducted to investigate the quantitative differentiation of FMs and calcification in each contrast. 3D U-Nets were developed using fastMONAI to automate the segmentation task using various combinations of MR-derived contrasts, with a model trained on CT used as a baseline. Models were evaluated using precision and recall calculated using a leave-one-out cross-validation scheme.

**Results:** Significant differences were observed between FM and calcification regions in CT, QSM and *T* 2^*^, though overlap was observed in QSM and *T* 2^*^. The baseline CT U-Net achieved an FM-level precision of ≈ 98% and perfect recall. The best-performing QSM-based model achieved precision and recall of 80% and 90%, respectively, while conventional MRI had values below 70% and 80%, respectively. The QSM-based model produced segmentations with good agreement with the ground truth, including a challenging FM that coincided with a bleed.

**Conclusion:** The model performance highlights the value of using QSM over indirect measures in MRI, such as signal voids in magnitude-based contrasts. The results also indicate that a U-Net can capture more information about the presentation of FMs and other sources than would be possible using susceptibility quantification alone, which may be less reliable due to the diverse presentation of sources across a patient population. In our study, QSM was a reliable discriminator of FMs and other sources in the prostate, facilitating an accurate and streamlined approach to MR-only RT.

## 1 Introduction

Prostate cancer (PCa) is the second most common cancer in men, with over 1.2 million new cases and 350,000 deaths annually [1]. For localized PCa, treatments include surgery and various forms of radiotherapy (RT), with intensity-modulated RT (IMRT) emerging as the standard of care, primarily due to its ability to target tumor volumes while minimizing doses to surrounding tissues precisely. However, geometric misses can occur due to inter- and intra-fractional patient motion [2, 3]. Such misses can be mitigated using gold fiducial markers (FMs) implanted into the prostate gland under ultrasound guidance for image-matching purposes [4, 5]. In the conventional CT-MRI planning workflow, MRI is used for its superior soft-tissue contrast in target delineation. Meanwhile, CT provides the attenuation information necessary for dosimetry and FM visualization. In the treatment room, the available imaging modality, typically kilovoltage or megavoltage X-ray, is used to verify the target positioning by matching the positions of FMs in the co-registered planning images. Recently, interest in MR-only RT planning has grown [6–8]. An MR-only workflow uses planning images derived solely from MRI rather than the conventional CT-MRI workflow. This approach offers several benefits, including reduced geometric uncertainty by eliminating the need for CT-MR registration, reduced cost, a more streamlined patient imaging pathway, and reduced patient exposure to ionizing radiation.

Despite the potential advantages of MR-only RT planning, the approach poses several challenges. Conventional MRI’s non-quantitative nature complicates dose calculations, an issue now addressed by synthetic CT (sCT) derived from MRI, which aligns closely with CT-based planning [9–12]. Geometric distortions in MRI, especially at air-tissue boundaries, affect spatial accuracy and remain a challenge [13, 14]. A major concern remains in the detection of FMs in MRI. In MRI, FMs appear as signal voids, complicating differentiation from other void sources such as calcifications and bleeds.

Since differentiating void sources in the prostate can be challenging and time-consuming, a range of automated FM detection techniques have recently been proposed for MR-only RT, with initial approaches based on template matching. An early method of this type used spectral clustering of FM templates constructed from manual contours, achieving FM detection accuracies of 63% and 88% for *T* 2^*^- and T1-weighted GRE sequences, respectively, over 15 patients [15]. Expanding on this technique, another study applied simulated complex-valued templates, reporting a true positive rate (TPR) of 96% across 17 patients [16]. Both investigations highlighted the challenges faced in distinguishing FMs from other sources such as calcifications and bleeds. Another method used logistic regression to integrate features across multi-parametric MRI and achieved a TPR of 94% over 32 patients [17]. A similar approach cross-referenced data from various MRI sequences to pinpoint FM candidates in T2w images, resulting in a TPR of 84% across 40 patients, with most false positives due to calcifications [18]. A significant breakthrough came with the adoption of deep learning techniques, where one study showcased a detection sensitivity of 97.4% across 39 prostate cancer patients, matching human expert performance in the same images [19].

Quantitative susceptibility mapping (QSM) is an MRI post-processing technique which calculates the magnetic susceptibility distribution of imaged objects from the static magnetic field [20–23]. Since QSM is sensitive to susceptibility variations, it has the potential to identify and differentiate FMs from calcification quantitatively. To date, QSM has mainly been applied in human brain imaging applications [24]. Investigations of QSM in the body present several challenges. First, the chemical shift effect of protons in fat can cause phase discontinuities and geometric distortions at fat-water interfaces [25], which gives rise to unwrapping and other artifacts in QSM. Second, the higher dynamic range of susceptibility values in body regions is problematic for regularizers optimized for narrower ranges in brain imaging. However, QSM investigations in the body are rapidly expanding with the development of advanced water-fat separation techniques, robust QSM reconstruction pipelines, and artifact reductions [26–31].

Initial work in prostate QSM investigated the visualization of post-biopsy calcification to aid in PCa diagnosis [32]. Realizing the potential for QSM to enable MR-only RT planning, later studies investigated QSM for localizing calcification as a natural alternative to implanted FMs [33], a concept initially applied in cone-beam CT [34]. Concurrent work specifically investigated using QSM to differentiate FMs from calcification *in silico* [35], including paramagnetic low dose-rate (LDR) brachytherapy seeds for post-implant dosimetry [36]. The same authors later expanded their work in LDR seeds to three patients for *in vivo* validation [37]. These LDR markers are paramagnetic and appear hyperintense in QSM, making them straightforward to differentiate from diamagnetic calcifications appearing hypointense. However, the gold FMs conventionally used for prostate IGRT are diamagnetic, and differentiation in QSM will rely more heavily on quantification or complementary MRI contrasts. Recent work has demonstrated that QSM may provide sufficient information for the differentiation task with a dataset of 7 PCa patients [38]. However, FM identification or differentiation requires validation using larger datasets and is yet to be automated.

This study investigates the effectiveness of a QSM post-processing pipeline optimized for body imaging applications in identifying and differentiating FMs from calcifications in a dataset of 26 PCa patients. Initially, the differentiation capability of multiple MRI contrasts, including QSM and *T* 2^*^ maps, is quantitatively assessed through statistical testing. Subsequently, 3D U-Nets are developed to automate the classification and segmentation of each region. The models are trained using combinations of MRI contrasts, including QSM, *R*2^*^, SWI, T1-weighted, and GRE magnitude images, to identify which combinations provide the most valuable information for the segmentation task. By mitigating the imaging challenges posed by calcifications in the presence of FMs, this work facilitates a more accurate and streamlined approach to MR-only RT.

## 2 Methods

### 2.1 Data and Acquisition

MRI and CT were acquired from 26 PCa patients, each with three implanted Gold Markers (Riverpoint Medical, Portland, Oregon, United States), cylindrical and 1 *×* 3 mm. All data were acquired with the approval of the relevant ethics committees. Given the study’s focus on differentiating FMs and calcifications, PCa patients with calcifications were preferentially scanned. In this sample, calcifications were present in 14 patients (≈ 54%).

CT data were acquired using a Siemens SOMATOM Confidence CT scanner (Siemens Healthcare, Erlangen, Germany) on software version VB10A (imaging parameters: imaging volume superior scan limit included L5 to lesser trochanters inferiorly, imaging resolution 0.98 *×* 0.98 *×* 2 mm, an image matrix size of 512 *×* 512 *×* 188, and kVp=120 kV).

MRI data were acquired using a MAGNETOM Skyra 3T scanner (Siemens Healthcare, Erlangen, Germany) with a 32-channel spine coil and 18-channel body coil on software version VE11. Acquisitions include T1-weighted anatomical scans and a gradient-echo (GRE) sequence used for QSM reconstruction. The GRE used a bipolar readout scheme and 1.4 mm^3^ isotropic resolution, TEs=2.46/4.92/7.39/9.84/12.3/14.8/17.22 ms, TR=25 ms, TA=2:54 min:sec, and FA=45°. Phased array channels were combined using the pre-scan normalize plus adaptive combine approach [39]. Due to the chemical-shift effect causing opposing geometric shifts between odd and even echoes, only the even, in-phase echoes were kept for further post-processing.

### 2.2 Data post-processing

GRE images were processed using QSMxT v5.1.0 [30, 40] using the default settings for body imaging applications. The body imaging pipeline applies TGV-QSM [41, 42] for the phase unwrapping, background field correction, and dipole inversion steps, an automated threshold-based masking algorithm, and a two-pass artifact-reduction algorithm. QSMxT also produced susceptibility-weighted imaging (SWI) and *T* 2^*^/*R*2^*^ maps using CLEAR-SWI [43] and NumART2star [44], respectively, from the MriResearchTools library [45].

CT images were registered and resampled to the GRE space using ITK-SNAP [46]. This resampling enabled a uniform U-Net training procedure by eliminating the need for architecture modifications across the modalities, ensuring a fair comparison of learning potential at the MRI resolution. The authors roughly segmented FMs, calcification, and prostate tissue regions separately for the registered CT and GRE images. To ensure a consistent segmentation approach, the rough segmentations were automatically filtered. The GRE FM segmentations were filtered to include susceptibilities three standard deviations below the mean prostate susceptibility, while calcifications were constrained by two standard deviations. In the CT images, FM segmentations were filtered to include values ten standard deviations above the prostate mean attenuation, while calcifications were constrained by four standard deviations. These thresholds were determined empirically, ensuring segmentations covered the distinct FM and calcification regions without including surrounding tissues. Segmentations were further refined using Gaussian smoothing and morphological operations.

The presentation of calcifications can vary considerably compared with FMs. Calcifications can be large or small, with varied levels of concentration. While the U-Net models were trained using all calcification segmentations, statistical analyses included only calcifications that were of similar size to FMs and which could be considered challenging FM candidates. This final filtering step included only calcifications within 3 standard deviations of the mean FM size in voxels.

### 2.3 Model and training

A 3D U-Net was implemented using fastMONAI [47, 48], which interfaces with fastai [49], MONAI [50], and PyTorch [51]. The U-Net was based on the fastMONAI example for multi-class segmentation, which comprises five levels with channels (16, 32, 64, 128, 256) and strides (2, 2, 2, 2), along with two residual units in each level. The U-Net was adapted to take multi-modal images as inputs and output predictions for background, calcification, and FM regions. Models were trained on various combinations of images to investigate how they impact performance. These combinations include standalone modalities of CT, QSM, SWI, T1, *R*2^*^, and GRE (combined across echoes using the sum-of-squares), as well as combinations of (QSM, T1, and *R*2^*^), (QSM and T1), and (QSM and SWI).

For training, input images and masks were initially cropped to 80 *×* 80 *×* 80 voxels, and data were processed in batches of six. The training was configured for 800 epochs, with early stopping triggered after 200 epochs of no improvement. The datasets were augmented using random flipping along the left-right and anterior-posterior axes, random affine transformations with rotations up to 90 degrees, and Z-normalization. To handle possible intraprostatic bleeds causing FMs to appear hyperintense in QSM, random multiplications of -1 were made to the FM QSM values in 5% of training examples. The model was optimized using the RAdam optimizer with the Lookahead method, known as Ranger, and an initial learning rate of 0.003 was set. A cosine annealing learning rate schedule was employed during the training process. The loss function combined Dice loss and cross-entropy loss with one-hot encoding applied to the target masks.

For evaluation, a leave-one-out cross-validation (LOOCV) scheme was implemented. This approach required training 26 models or *folds*, each with a different validation example. This approach allows for a more reliable assessment of model performance without sacrificing valuable training examples for a separate testing set. No hyperparameter tuning was conducted to minimize the risk of overfitting to the validation examples. Each model’s performance was evaluated by averaging metrics across the 26 folds. Metrics for evaluation are the multi-class Dice score, voxel-level true positive and false positive rates, voxel-level recall and precision, and FM-level recall and precision. The multi-class Dice score quantifies the overlap between predicted and ground truth segmentations. Precision and recall reflect the model’s ability to correctly identify FMs, differentiate them from other sources, and recall actual FMs reliably. Voxel-level metrics are calculated for individual voxels, whereas FM-level metrics are calculated across whole FMs.

## 3 Results

78 FMs were observed across each contrast and modality, corresponding to the expected 3 FMs per patient. FMs appeared hyperintense in CT and hypointense in T1-weighted, SWI, and GRE magnitude images. In QSM, 75 FMs appeared hypointense, one appeared hyperintense, and the remaining two appeared isointense (see Figure 1). A radiologist confirmed the hyperintensity as a bleed at the FM insertion site. The two isointensities occurred in one patient and coincided with hyperplasic tissue due to benign prostatic hyperplasia (BPH) and a possible bleed. The *R*2^*^ images were noisy, and FMs were difficult to discern but tended towards hyperintense compared to the surrounding prostate tissue.

**Figure 1:**
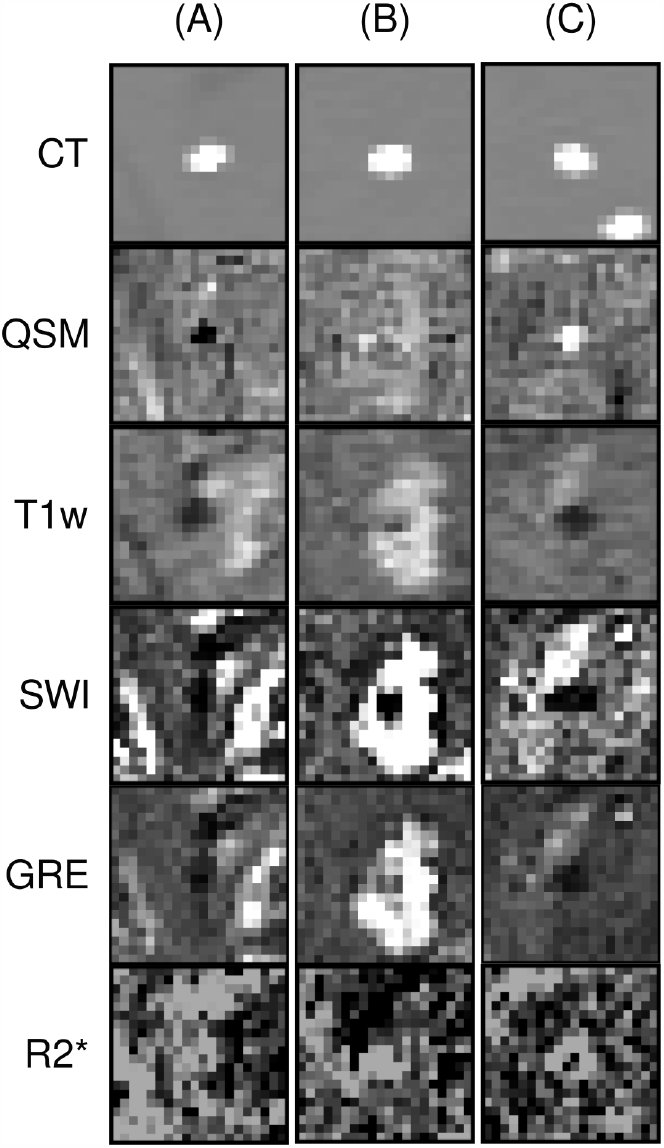
Presentation of FMs across image types; cropped axial regions. A) depicts a typical FM appearing hypointense in QSM; B) depicts an FM appearing within hyperplasic tissue and a small bleed which appears isointense in QSM; C) depicts an FM coinciding with a bleed which appears hyperintense in QSM. The typical FM appearance in QSM has a clearer boundary than in the other MR-derived images.

Statistical significance was observed across the distributions of mean FM and calcification values in the quantitative images, including CT, QSM, and *T* 2^*^ (see Figure 2). There was no overlap between the calcification and FM mean value distributions in CT histograms, though overlap was observed in QSM, with more extensive overlap in *T* 2^*^.

**Figure 2:**
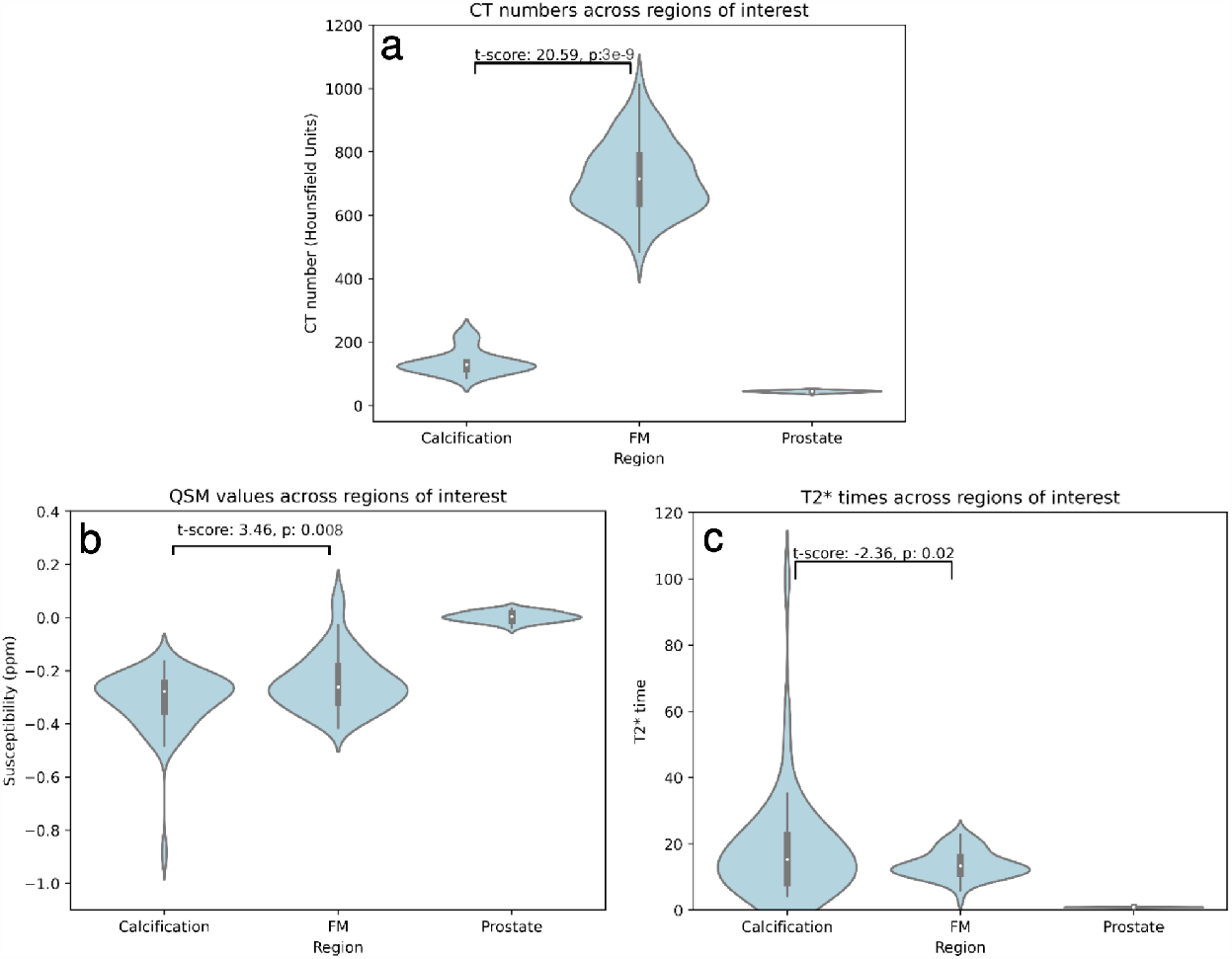
Region-level mean values for calcifications, FMs, and prostate tissue in A) CT numbers; B) QSM values; and C) *T* 2^*^ times. Calcifications were filtered to those with sizes within three standard deviations of the mean FM size. Violin widths are equal. The t-scores and associated p-values for the calcification and FM regions are annotated. While there is significant differentiation between the means of the FM and calcification distributions in all contrasts, there is some overlap in QSM and *T* 2^*^.

The performance of each U-Net was evaluated using Receiver Operating Characteristic (ROC) and Precision-Recall (PR) curves. Curves were computed based on the FM labeling of individual voxels, with the metrics averaged across cross-validation folds. For the ROC curves (see Figure 3), CT had the highest performance with an area-under-curve (AUC) of 0.99, followed closely by the QSM model (AUC of 0.97). More conventional MRI trailed behind the models incorporating QSM, with AUCs ≤ 0.87. CT again demonstrated superior performance in the PR curves with AUC = 0.91 (see Figure 4). Models incorporating QSM were clustered with AUC ≈ 0.65, with other MRI formed a separate cluster with inferior AUCs ≈ 0.3.

**Figure 3:**
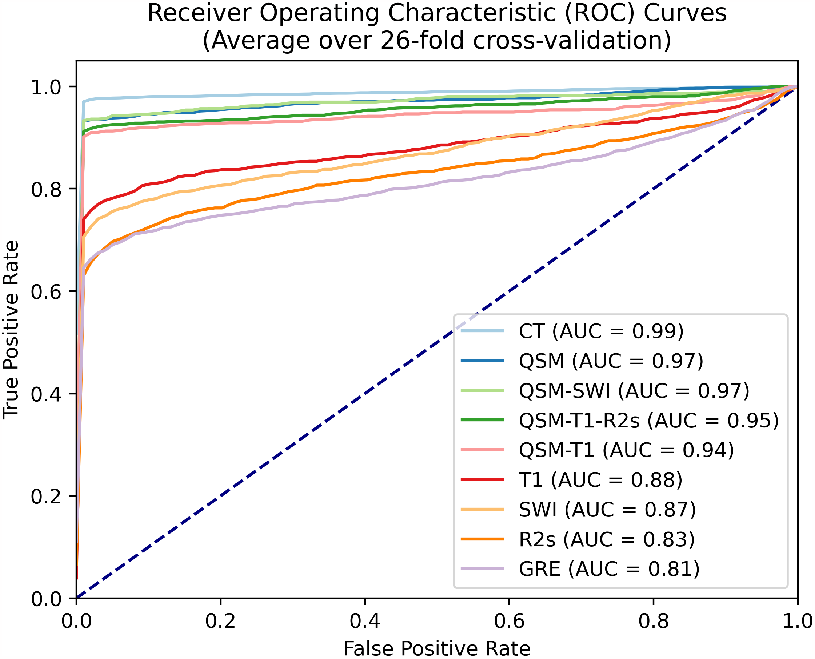
Voxel-level Receiver Operating Characteristic (ROC) curves for each imaging modality on validation data and evaluated against the FM label, averaged over all cross-validation folds.

**Figure 4:**
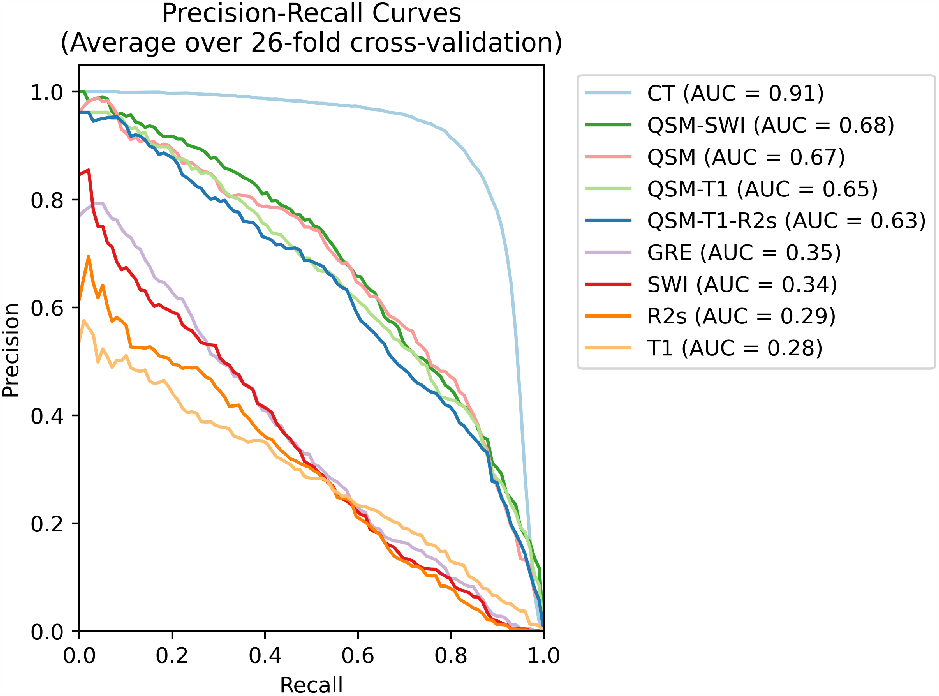
Voxel-level Precision-Recall (PR) curves for each imaging modality on validation data and evaluated against the FM label, averaged over all cross-validation folds.

Precision and recall metrics were also computed at the FM level across models and averaged across cross-validation folds (see Table 1). The CT model achieved a precision of 0.98 *±* 0.07 and a perfect recall. The QSM model achieved a precision of 0.8 *±* 0.21 and recall 0.9 *±* 0.2, similar to the other models incorporating QSM. Models trained using other MR-derived images, including SWI, T1, *R*2^*^ maps, and GRE magnitude images, had achieved precision ≤ 0.67 and recall ≤ 0.79, along with wider standard deviations.

**Table 1:** FM-level precision and recall across models, averaged over cross-validation folds.

| Model | Precision | Recall |
| --- | --- | --- |
| CT | $0.98 \pm 0.07$ | $1 \pm 0$ |
| QSM-T1-R2s | $0.81 \pm 0.25$ | $0.9 \pm 0.25$ |
| QSM-T1 | $0.77 \pm 0.25$ | $0.87 \pm 0.28$ |
| QSM-SWI | $0.8 \pm 0.2$ | $0.93 \pm 0.19$ |
| QSM | $0.8 \pm 0.21$ | $0.9 \pm 0.2$ |
| T1 | $0.67 \pm 0.31$ | $0.79 \pm 0.34$ |
| SWI | $0.57 \pm 0.28$ | $0.65 \pm 0.35$ |
| R2s | $0.61 \pm 0.27$ | $0.71 \pm 0.31$ |
| GRE | $0.56 \pm 0.31$ | $0.59 \pm 0.36$ |

Representative QSM images depicting FMs and calcification were visualized along with ground truth segmentations, model predictions using the QSM U-Net, and resultant labels (see Figure 5). The model’s predictions are closely aligned with the ground truth labels.

**Figure 5:**
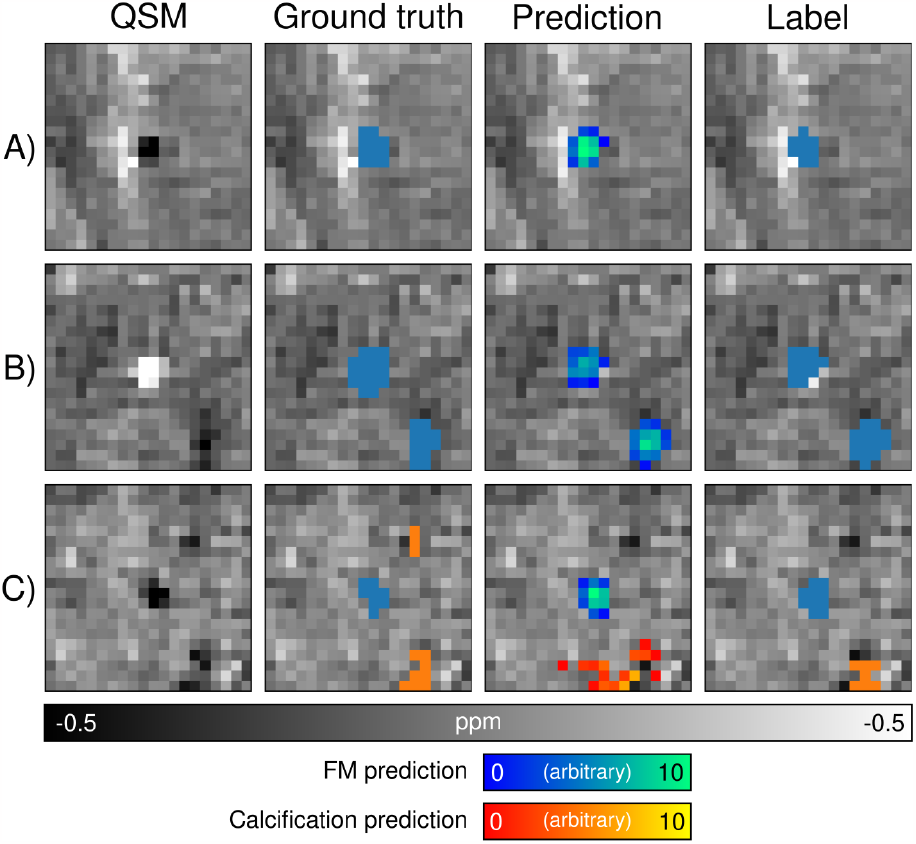
Representative examples of FMs and calcification in QSM, including ground truth segmentation labels, predictions produced by the segmentation model trained on QSM only (in arbitrary units), and the predicted label based on the maximum prediction across each class. A) depicts a typical FM; B) depicts an FM appearing hyperintense due to a bleed at the insertion site; C) depicts an FM with nearby calcification.

## 4 Discussion

This study evaluated the efficacy of a QSM post-processing pipeline for distinguishing intraprostatic FMs from calcifications in 26 PCa patients in conjunction with various MRI contrasts and automated the segmentation task with a 3D U-Net, facilitating a streamlined MR-only RT approach.

Our initial statistical analysis showed significant differences in mean CT numbers, QSM values, and *T* 2^*^ times of distinct FM and calcifications. While there was no overlap between these regions in CT numbers, the QSM values and *T* 2^*^ times had some overlap. This indicates that discriminating the sources solely on MR-based quantitative values remains challenging despite the significant differences in their distributions.

U-Nets were trained for FM and calcification segmentation using single or multi-contrast input. The standalone CT model excelled in identifying FMs with 98% precision and perfect recall. Meanwhile, QSM-based models showed commendable performance, with the standalone QSM model achieving 80% precision and 90% recall. In contrast, models trained on MRI without QSM performed poorly, with precision and recall values below 70% and 80%, respectively.

Prior work investigating FM localization using QSM demonstrated quantitative differentiation of FMs from calcifications [38]. The authors measured FMs at -31.5 *±* 2.0 ppm and calcifications at -14.6 *±* 0.9 ppm in a dataset of seven PCa patients, two of which were reported to have calcifications. Our study did not replicate these findings, with extensive overlap in mean susceptibility values between FMs and calcifications across 26 PCa patients. However, a direct comparison is challenging because of the small number of calcifications in the cited work and a lack of clarity on FM and calcification segmentation methods. Moreover, calcification presentation can vary considerably in size and concentration, and the material and size of the FMs used could have a substantial influence on findings. Our analysis considered the mean values of distinct calcified regions, similar in size to the 1 *×* 3 mm pure gold FMs, segmented semi-automatically as described in the methods section.

Most automated FM detection techniques in existing literature rely on magnitude images [15, 18, 19], with some including multi-parametric MRI [17, 18], and another using complex-domain images [16]. The visualization of FMs as distinct signal voids arises from their underlying magnetic susceptibility difference from prostate tissues. This insight motivated our investigation into techniques based on QSM images, which offer a more direct susceptibility measurement. Although the best-performing technique to date uses magnitude images alone [19], it is worth noting that the cited study used larger FMs than the present study, which could explain their detection efficacy. In the present study, several U-Nets were trained using the multi-parametric approach with combinations of MR images including (QSM, T1, and *R*2^*^), (QSM and T1), and (QSM and SWI) to determine whether they could enhance a deep-learning-based solution. As hypothesized, the U-Nets trained on QSM images surpassed the performance of those trained on T1, R2, SWI, and GRE magnitude images. However, combining multiple contrasts did not markedly improve performance in our study. In some instances, performance even deteriorated slightly, possibly due to the introduction of noise or redundant information that detracted from the distinguishing features offered by QSM alone.

The performance of the U-Nets may be improved further through hyperparameter tuning. Avoiding this in our study was a deliberate choice to prevent overfitting to the limited dataset. The LOOCV approach facilitates a fair evaluation of the U-Nets and maximizes training data availability while reserving only one example for evaluation in each of the 26 folds. However, a more extensive dataset may facilitate a separate testing set and permit hyperparameter tuning to optimize performance on validation data before final testing. Another reason hyperparameter tuning was avoided was to ensure the same network architecture could be applied for each U-Net regardless of the input image combinations. However, each image type or combination might require individualized hyperparameter adjustments to perform optimally.

Another way the performance of the U-Nets may be improved is by tweaking the acquisition. The current best-performing FM detection technique used an acquisition with a slice thickness of 2.8 mm and an in-plane resolution of 1.46 mm [19], gathering more signal per voxel as in our isotropic resolution of 1.4 mm^3^. The isotropic voxel size was chosen because it improves quantification in QSM as recommended in the recent QSM consensus paper [52]. Investigating sequence optimizations, such as altering the slice thickness, may improve model performance and visualization of sources in QSM for this particular application.

Multiple studies have noted that differentiation of FMs from other sources, such as bleeds and calcification, can be challenging in magnitude images [15, 16, 18]. This generally occurs with calcifications that result in a signal void similar in size and shape to those caused by FMs. Previously proposed automated methods have resulted in false negatives where bleeds occur at the FM insertion site. Although our dataset contains only a single instance of such a bleed, the data augmentations that simulate bleeds aided the QSM U-Net in identifying the FM successfully. This outcome highlights the benefits of realistic simulated datasets and augmentations, which are becoming increasingly important for clinical deep-learning tools where large-scale datasets are challenging to acquire [53, 54].

One major limitation of the present study is the dataset’s relatively small size. While our dataset of 26 PCa patients is more extensive than similar studies, it may not capture the anatomical variability of a broader population. Uncommon cases were scarce, such as confirmed bleeds at FM insertion sites, instances of BPH, or other gross pathology causing complex tissue composition changes. This paucity could affect the U-Net’s ability to identify FMs at clinically essential edge cases effectively. Therefore, in addition to realistic simulations and data augmentations, acquiring larger datasets over broad populations will be crucial to understanding these edge cases and developing more robust models.

Future research should aim to build larger datasets encompassing a more diverse cohort of PCa patients. This expansion would improve model performance, generalizability, and robustness across patient subsets. The current findings in uncommon cases like bleeds emphasize the importance of obtaining more such data for richer insights into their QSM and conventional MRI interpretations. While our simulated bleeds approach proved effective, there is potential to leverage advanced augmentation techniques further. As findings evolve, assessing the practical utility of automated FM detection tools in clinical contexts will become crucial, comparing their outcomes with manual radiologist annotations to determine clinical readiness.

## 5 Conclusions

This study evaluates the potential of QSM to identify and distinguish intraprostatic FMs from calcifications in 26 PCa patients. The developed U-Nets subsequently automate the segmentation task. The model performance highlights the value of using QSM over indirect measures such as signal voids in magnitude-based contrasts and SWI. The results also indicate that a U-Net can capture more information from the presentation of FMs and other sources than would be possible using susceptibility quantification alone, which may be less reliable due to the diverse presentation of sources across a patient population. In our study, QSM was a reliable discriminator of FMs and other sources in the prostate, facilitating an accurate and streamlined approach to MR-only RT. Future work should focus on expanding training datasets to capture the variability across wider patient populations and develop clinical tools to automate the FM contouring and localization task.

## 6 Data Availability Statement

All code used is publicly available on GitHub. QSMxT for QSM reconstruction: https://qsmxt.github.io/ (v5.1.0). U-Net architecture and training code: https://github.com/astewartau/prostate.

## 8 Acknowledgments

We thank the patients involved in this study and the radiation oncology team at Calvary Mater, Newcastle, Australia. The authors SB and AWS acknowledge funding through an ARC Linkage grant (LP200301393) and acknowledge support through the UQ AI Collaboratory. MB acknowledges funding from the Australian Research Council Future Fellowship grant FT140100865 and SR from the Austrian Science Fund (FWF): 31452. This research was partly conducted by the Australian Research Council Training Centre for Innovation in Biomedical Imaging Technology (project number IC170100035) and funded by the Australian Government. The authors would like to acknowledge Siemens Healthineers for supporting our project by providing the WIP sequence used to acquire the QSM data in this publication. This research was undertaken with the assistance of resources and services from the Queensland Cyber Infrastructure Foundation (QCIF) and the UQ Research Computing Centre (RCC).

